# Comparative analysis of the effect of genomic isolators flanking transgenes to avoid positional effects in Arabidopsis

**DOI:** 10.1101/384487

**Authors:** Ana Pérez-González, Elena Caro

## Abstract

**Highlight:** We have studied the effect of different insulator sequences over transgene expression levels and variability, and over transgene integration, using NGS. Our results compare the benefits obtained by their use.

**Abstract:** For more than 20 years, plant biologists have tried to achieve complete control of transgene expression, but until gene targeting techniques become routine, flanking transgenes with genetic insulators can help avoid positional effects. Insulators are DNA sequences with barrier activity that protect transgenes from interferences with the host genome. We have, for the first time, compared the effect of three insulator sequences previously described in the literature and of a matrix attachment region from Arabidopsis never tested before. Our results indicate that the use of all sequences increases transgene expression, but only the last one reduces variability between lines and between individuals to a minimum. We have analyzed the integration of insulator-flanked T-DNAs using whole genome re-sequencing (to our knowledge, also the first time) and found chiMAR lines with insertions located within heterochromatic regions of the genome, characterized by DNA methylation that did not spread into the T-DNA, suggesting that chiMAR can shelter transgene insertions from neighboring repressive epigenetic states. Finally, we could also observe a loss of accuracy of the RB insertion in the lines harboring insulators, evidenced by a high frequency of truncation of T-DNAs and of insertion of vector backbone that, however, did not affect transgene expression.

## Introduction

Due to the random nature of transgene insertion in the majority of higher eukaryotes, transgenic DNA may integrate into regions of the genome that are transcriptionally repressed (heterochromatin), which can result in many cases in transgene silencing. Additionally, transgenes may be incorporated near endogenous regulatory elements, such as transcriptional enhancers or repressors, which can cause their miss-expression (reviewed by Pérez-González & Caro 2016).

Chromatin insulator sequences, or boundary elements, are DNA sequences with the capacity to define a chromatin domain because of two key activities, the first is the ability to interfere with enhancer-promoter communication when placed between the two (enhancer blocking activity) and the second one is the ability to protect a flanked transgene from position-dependent silencing (barrier activity) (Matzat and Lei, 2014).

These barrier elements have been characterized extensively in animals. In plants, possibly the best studied elements with potential applications are scaffold or matrix attachment regions (S/MARs), which have been suggested to trigger the formation of chromatin loops, and thus delimit the boundaries of discrete chromosomal domains (Butaye et al., 2004). Much of the research carried out concerning the use of transgene-flanking MARs as genetic insulators has shown that the use of these elements results in an increase in the level of transgene expression and/or a reduction in plant-to-plant variability (Butaye et al., 2005). However, in some cases, reports of success using this technique have been followed by negative results (De Bolle *et al*., 2003, De Bolle *et al*., 2007).

One of the most studied MARs is the one localized upstream the chicken lysozyme gene (chiMAR) (Loc and Strätling, 1988). Its role as insulator was shown in studies with animal cell lines where its presence near a reporter gene produced an increase in transgene expression and a decrease in variability among different lines (Stief et al., 1989). The use of the chiMAR in plant constructs has been somehow controversial, leading to reports with different conclusions. Mlynarova *et al*., 1994 showed that the chicken sequence was able to bind to the tobacco nuclear matrix and that when it flanked a T-DNA containing a GUS reporter gene, the variability of its expression decreased in full-grown primary transformants of tobacco. The same group later found that a significant reduction in variation of gene expression was conferred upon the GUS gene driven by the double cauliflower mosaic virus 35s promoter, but not to the NPTII gene, driven by the nopaline synthase (pNOS) promoter (Mlynarova et al., 1995). These results could, however, not be replicated in *Arabidopsis thaliana* first generation plants, where the chiMAR was found to have no influence on the level or variability of expression of transgenes driven by the 35S promoter (De Bolle et al., 2003). In fact, later studies applying different transformation methods and plant species reported no boost effect on transgene expression of wild type plants (De Bolle et al., 2007), but an increase in silencing mutant backgrounds (Butaye et al., 2004).

Allen *et al*., 1996 showed that stably transformed cell lines in which a GUS reporter gene was flanked by the tobacco MAR isolated from a genomic clone containing a root specific gene (Rb7) (Hall et al., 1991) produced more than 140 times more GUS enzyme activity than control transformants without it. However, the use of Rb7 did not reduce variation between different transformants.

The effect of the Rb7 MAR increasing transgene expression was also reported by Mankin *et al*., 2003, that analyzed in depth the specificity of the results depending on the promoter used. They reported that highly active promoters exhibited significant increases in GUS activity in constructs flanked by Rb7 compared to controls, but its presence did not significantly increase GUS activity when driven by weak promoters. Importantly, most transgenes flanked by the insulator showed a large reduction in the number of low expressing GUS transformants, suggesting that MARs can reduce the frequency of gene silencing.

Following that line, Abranches *et al*., 2005 tested the effects of Rb7 in conjunction with regulated transcription using a doxycycline-inducible luciferase transgene. The Rb7 lines showed higher reporter gene expression levels and avoided silencing apparition in the absence of active transcription from condensed chromatin spreading.

Another well characterized genetic insulator, defined initially by its ability to block interactions between enhancers and promoters when positioned between them, is the petunia transformation boost sequence (TBS) (Hily et al., 2009). This sequence has been shown to function in Arabidopsis and tobacco, and a detailed analysis of the motifs it contains showed that several specific regions are required for maximum enhancer-blocking function (Singer et al., 2011).

It was only a few years ago that another work showed that the TBS could similarly function in synthetic constructs sheltering transgenes promoters from the host plant genome regulatory elements. The TBS sequence was found to produce enhanced transgene expression, but did not prevent gene silencing in transformants with multiple and rearranged gene copies (Dietz-Pfeilstetter et al., 2016).

Almost 25 years after the description of some of these DNA sequences, their use is still not common practice in plant engineering projects due to their big size that makes troublesome cloning them through traditional methods, and because the reports on their effect are scattered over different organisms and transformation methods with no comparisons to allow for comparison between them.

Targeting transgenes to a specific integration site in the plant genome might rule out chromosomal position effects, but until there are routine efficient techniques for plant directed gene targeting, another alternative method needs to be developed.

With the advent of modular cloning techniques that allow rapid and straight forward generation of multigene constructs, the incorporation of genetic insulators to the flanks of T-DNAs is no longer a problem. Therefore, we decided to perform a systematic and parallel study comparing the activity and effectivity of incorporating different boundary elements flanking transgenes as a strategy in T-DNA design to maximize and stabilize transgene expression. We have, moreover, used whole genome re-sequencing for the molecular characterization of the insertion of insulator-flanked T-DNAs, finding interesting results that point to previously unknown functions of the barrier sequences.

## Material and Methods

### Modular cloning

Modular pieces AtS/MAR10 and Rb7 were amplified by PCR using Phusion High-Fidelity DNA Polymerase (NEB) from *A. thaliana* and *N. tabacum* genomic DNA using primers 359/348 and 269/270, respectively; chiMARs and TBS were amplified using KAPA2G Fast HotStart DNA Polymerase (Sigma) from chicken liver tissue and *P. hybrida* genomic DNA using primers 1724/1725/1726/1727 and 275/276/277/278, respectively.

Modular pieces were cloned into pFranki (chiMARs and TBS) or into GoldenBraid pUPD2 (Rb7 and AtS/MAR10) vectors, as described in (Sarrion-Perdigones et al., 2011). pFranki is a home-made vector adapted to clone pieces originally designed for GB2.0 so they can be compatible with GB3.0 and MoClo cloning systems. pFranki vector is composed by the cloning cassette of the GoldenBraid pUPD vector and the backbone of the pUPD2 vector. To generate transcriptional units, MoClo Level1 destination vectors were used (pICH47732-L1 P1, pICH47742-L1 P2, pICH47751-L1 P3, pICH47761-L1P4). Insulators modular pieces were cloned into L1P1 and L1P4 in all cases. Luciferase transcriptional unit was cloned into L1P2 vector using the following modular pieces: pICH85281 (pMAS), pICSL80001 (luciferase CDS), pICH41421 (tNOS) (Engler et al., 2014). Bialaphos resistance cassette (pICSL70005) was cloned into L1P3. Level2 destination vector pAGM4673 (Weber et al., 2011) was used for multigene assembly, and a rule of 2:1 molar ratio of inserts:acceptor was applied for adding Level1 plasmids to the reaction. Level1 and Level2 digestion/ligation reactions were performed in a thermocycler as follows: 20 seconds at 37%C, [3 minutes at 37%C, 4 minutes at 16%C] for 26 cycles, 5 minutes at 50°C, 5 minutes at 80%C, hold 16%C (adapted from Weber et al. 2011). *E. coli* DH5α quimiocompetent cells were transformed with the ligation products from either level and grown in LB medium containing X-Gal (20μg/mL) (Duchefa) and IPTG (1mM) (Anatrace), supplemented with ampicillin (100μg/mL) (Formedium) for GB pUPD and MoClo Level1, chloramphenicol (50μg/mL) (Formedium) for GB pUPD2, or kanamycin (50μg/mL) (IBIAN Technologies) for MoClo Level2. Sequencing (Macrogen) was done previously to plant transformation for correct sequence confirmation.

### Plant transformation

Level 2 transformation plasmids were introduced into *Agrobacterium tumefaciens* LBA4404 quimiocompetent cells and plated in LB medium supplemented with Rifampicin (25μg/mL) (Sigma-Aldrich), Streptomycin (100μg/mL) (sigma-Aldrich) and Kanamycin (50μg/ml). A single transformant colony was grown in 200mL LB medium supplemented with the same antibiotics at 28%C under constant shaking to perform Col0 plant transformation (Clough and Bent, 1998).

### Plant growth conditions and selection

T1 seeds were put into soil and grown in an environment controlled room (FitoClima HP, Aralab) under 16/8 hours light/dark conditions, at 22%C and 65% RH. After 10-20 days, seedlings were sprayed with Basta herbicide (200mg/L). Resistant plants were grown in the same conditions for T2 seeds recovering.

Seedlings were grown in plates in MS medium (Murashige and Skoog, 1962) with 1% sucrose, supplemented with 6μg/mL of DL-Phosphinothricin (Basta) herbicide (DL-Phosphinothricin, Sigma-Aldrich) for selection when needed, in a growth chamber under 16/8 hours light/dark conditions at 22%C.

### Luciferase reporter assay

For luciferase imaging, 16 seedlings per line were sowed in plates to analyze LUC activity. D-Luciferin Firefly, potassium salt (Biosynth) was dissolved in sterile H2O with 0.01% Triton X-100 to a final concentration of 0.2μM and sprayed over. After 6 minutes in the dark, luciferase activity was measured in a NightOWL II LB 983 (Berthold Technologies), with 3 minutes of exposition.

### Whole Genome Re-sequencing

Isolation of Arabidopsis genomic DNA was performed using a DNeasy Plant Mini Kit (Qiagen). Samples were sent to Novogene Co., Ltd. for library construction and sequencing. There, genomic DNA of each sample was randomly sheared into short fragments of about 350bp. These fragments were subjected to library construction using the Illumina TruSeq Library Construction Kit, strictly following manufacturer’s instructions. As followed by end-repairing, dA-tailing and further ligation with Illumina adapters, the requirement fragments (between 300bp and 500bp) were selected by PCR and amplified. After gel electrophoresis and subsequent purification, the required fragments were obtained for library construction.

Quality control of the constructed libraries were performed afterwards. Qubit 2.0 fluorometer (Life Technologies) was used to determine the concentration of the DNA libraries. After that, a dilution to 1 ng/μl was done and the Agilent 2100 bioanalyzer was used to assess the insert size. Finally, a quantitative real-time PCR (qPCR) was performed to detect the effective concentration of each library. Pair-end sequencing was performed on the Illumina platform, with the read length of 150bp at each end.

### Bisulfite conversion and sequencing

Genomic DNA of 12 days-old plants of line chiMARs 6.13 was extracted using a DNeasy Plant Mini Kit (Qiagen). Bisulfite treatment was done using the EZ DNA Methylation Gold kit (Zymo Research) following the manufacturer’s instructions. Amplification from converted DNA was performed with NXT Taq PCR kit (EURx) using primers 642 and 635. PCR fragments were checked on an 1% agarose gel for size verification. 4μl of PCR product was cloned into pGEM-T Easy (Promega) and transformed into chemically competent *E. coli* DH5α cells. Nine clones were selected for the analysis. Plasmid DNA of each clone was sent for sequencing (GATC), and results were checked using Geneious version 10.2.2 software (Kearse et al., 2012). Comparison of the converted clones to the original unconverted sequences was done using CyMate software (Hetzl et al., 2007), to count the converted/unconverted cytosines at each site. Percenatge of methylation was calculated as (number of methylated C residues in each context (CG, CHG or CHH)/total number of C residues in that context)*100.

## Results

Since the advent of plant genetic transformation, plant biologists have tried to maximize transgene expression level and minimize variability by flanking transgenes with genetic insulators. There are numerous studies that describe the use of a certain insulator sequence in a host organism and analyze different aspects of its barrier and enhancer-blocking ability, but they are performed in such diverse conditions that do not allow for comparison and their results are sometimes contradictory. Our work consists on the use four different insulator sequences flanking a LUC transgene with the aim of conducting a definitive parallel and systematic analysis of their effect on transgene integration, expression level and variance in Arabidopsis.

Taking advantage of the capacities of modular cloning systems, we generated five identical constructs harboring the firefly luciferase transgene driven by the constitutive mannopine synthase Agrobacterium gene promoter (pMAS) and followed by the Basta resistance selection marker cassette. One of these constructs was used as a control, and the other four were flanked by different sequences reported in the literature to have some type of insulator activity (Figure 1A). The insulator sequences used in this work were the MAR located next to the tobacco root specific gene Rb7 (Rb7) (Hall et al., 1991), the chicken lysozyme A MAR region (chiMAR) (Loc and Strätling, 1988), the petunia transformation booster sequence (TBS) (Hily et al., 2009) and one of the scaffold/matrix attachment region sequences isolated from Arabidopsis chromosome 4 (AtS/MAR10) (Pascuzzi et al., 2014).

**Figure 1.**
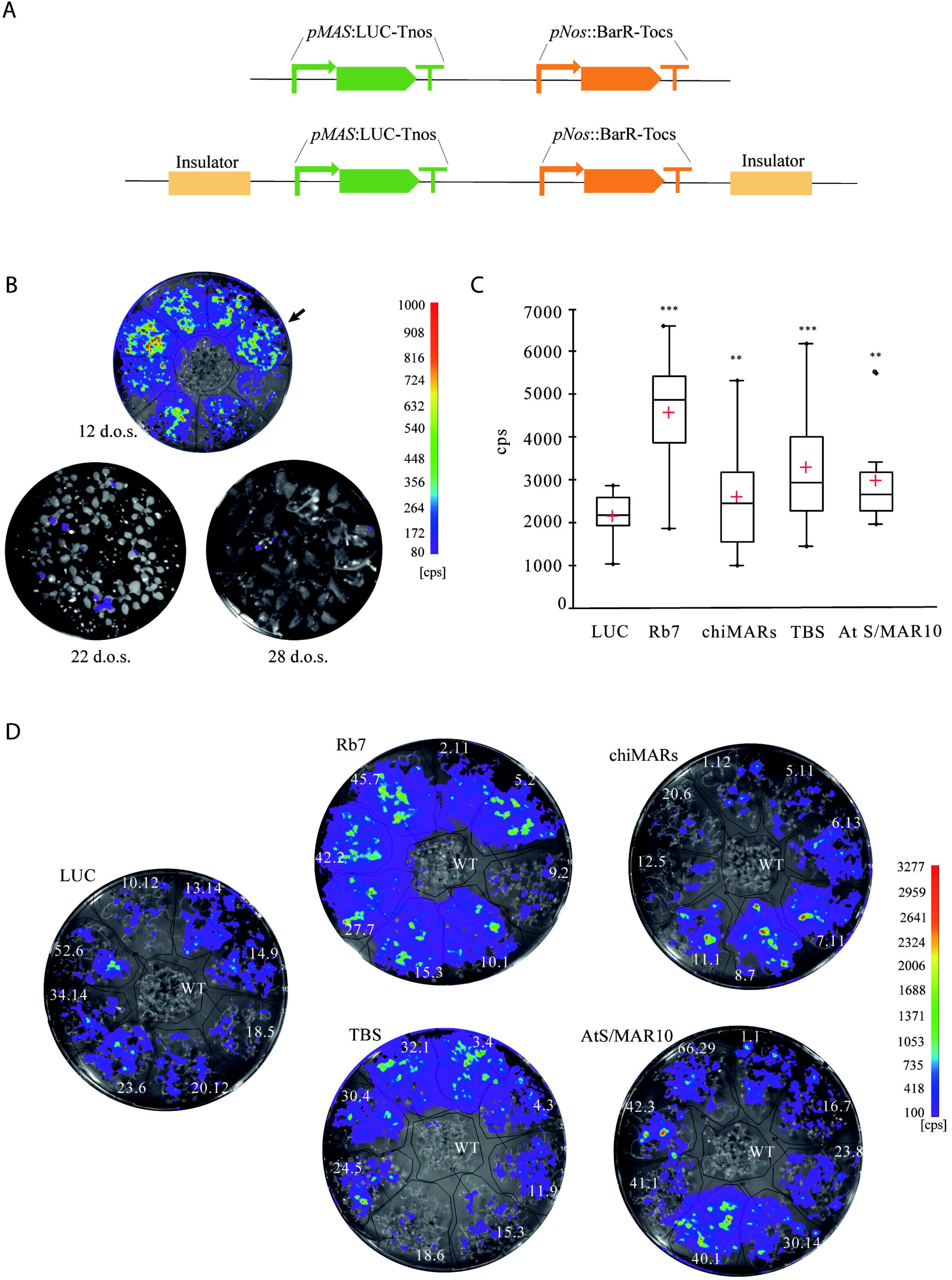
Analysis of insulator effect over LUC activity. A) Schematic representation of the constructs used for studying the effect genomic insulators flanking transgenes. The above scheme represents the construction used as a control (LUC) while the scheme below represents the four constructions flanked by the four different insulators. *pMAS:* mannopine synthase gene promoter; LUC: firefly luciferase; Tnos: nopaline synthase terminator: *pNos:* nopaline synthase promoter; Tocs: octopine synthase terminator. “Insulator” represents Rb7, chiMAR, TBS or AtS/MAR10. B) Time course of LUC activity when expressed under the *pMAS* promoter. Lines were assayed for LUC imaging at 12, 22 and 28 days-old. Results for control line LUC 14.9 (indicated with an arrow) are shown, but similar data was obtained for the rest of the lines. d.o.s: day-old seedlings; cps: counts per second. C) Box plots showing LUC activity. ** represents Student’s test significant differences (p<0.005); ***represents Student’s test highly significant differences (p<0.001); cps: counts per second. D) LUC activity imaging of the T3 homozygous lines, eight lines per construction.

A time course study of the LUC expression conferred by the pMAS showed that its activity was maximum in young seedlings, and decreased rapidly as plants matured and formed the rosette (Figure 1B). Given these results, for the following experiments, LUC activity was always measured in 12 day old seedlings. Eight 3:1 segregating Arabidopsis Col0 T2 lines were randomly selected and a 100% Basta resistant T3 line coming from each of them was used for LUC activity imaging to assess their levels of transgene expression (Figure 1C). Our results confirmed previous reports, indicating that all constructs flanked by insulator elements led to plants with increased transgene expression (Figure 1D).

Another property of insulator sequences is their ability to decrease variability between transgenic lines transformed with the same construct. When the transgene was flanked by Rb7, chiMAR or TBS, the increase in LUC expression described above was accompanied also by a statistically significant increase in the coefficient of variation between lines, which measures the extent of variation in relation to the mean within a population (Figures 2A and B). Line 40.01 from AtS/MAR10 behaved very differently from the rest in terms of expression (Figure 1B). We confirmed it was an outlier (expression value above Q3 + 1.5×InterQuartileRange) and thus, did not consider it for this analysis. When the outlier line data was removed, the presence of AtS/MAR10 flanking the transgene led to the opposite effect than the rest of insulators, a statistically significant reduction in the coefficient of variation between lines, or what is the same, a reduction in inter-line variation (Figures 2A and B).

**Figure 2.**
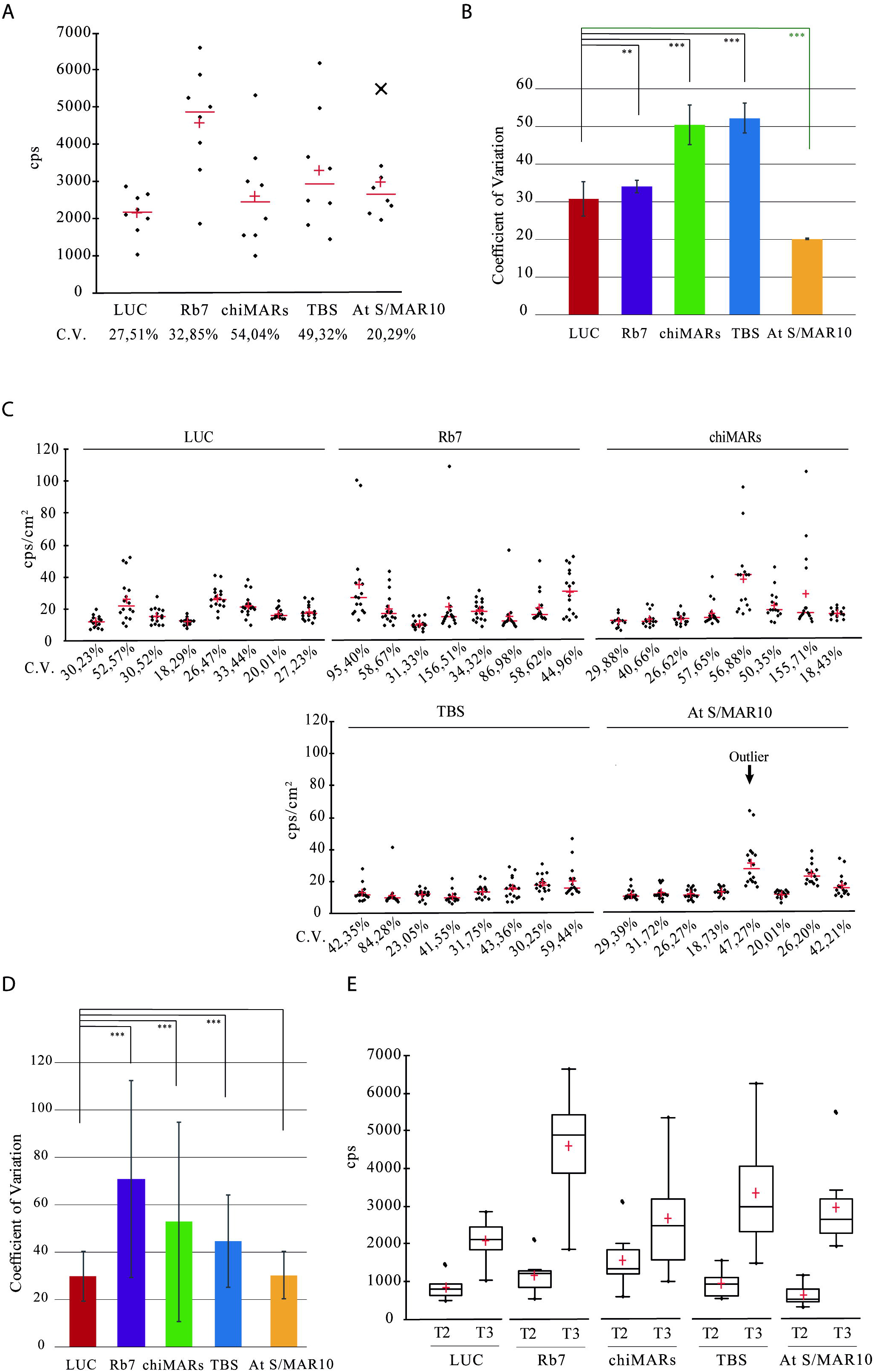
Analysis of insulator effect over inter-line, inter-individual and inter-generation variation of LUC activity. A) Scattergrams showing LUC activity in the selected eight lines obtained after transformation with each construct. The CV of each population was calculated as (standard deviation/mean)*100. B) Comparison of the inter-line coefficient of variation. ** represents Student’s test significant differences (p<0.005); *** represents Student’s test highly significant differences (p<0.001); CV: coefficient of variation; cps: counts per second. C) Scattergrams showing LUC activity in 16 seedlings of the eight selected lines obtained after transformation with each construct. CV was determined for each line and calculated as (standard deviation/mean)*100. cps/cm2: counts per second/cm2. The arrow in the AtS/MAR10 graph represents the outlier line. D) Comparison of the inter-individual coefficients of variation. CV for each insulator was calculated as (standard deviation/mean)*100. A great variance was overserved for the insulated lines compared to the control except for AtS/MAR10, that showed a small variation similar to the control, in agreement with the Student’s test. ***represents highly significant differences (p<0.001). E) Box plots showing LUC activity in T2 and T3 generations of the 8 selected lines obtained after transformation eighth each construct. cps: counts per second.

To measure the level of variation between genetically identical individuals within a population, we measured the expression of 16 seedlings from each line, and analyzed the effect of insulators on inter-individual (intra-line) variation (Figure 2C). For Rb7, chiMAR and TBS, the increase in expression induced was not homogeneous between individuals and, as a result, there was a greater variance in these lines compared to the control. For AtS/MAR10, there was a small variance, similar to that of the control with no insulator (CV around 25%) (Figure 2D).

Next, we compared LUC expression in segregating lines from the T2 generation with homozygous lines from the T3 generation, in an effort to establish if, in our system in study, LUC expression was dependent on gene dosage. Our experiments confirm an increase in expression in all T3 lines compared to T2, consistent with the establishment of homozygous populations. No differences could be observed due to the presence of insulators (Figure 2E).

In an effort to further characterize the insulators lines in more detail than previous works, we proceeded to perform whole genome re-sequencing (WGR) in some of the lines obtained by transformation with each construct (Figure 3A). The results allowed us to select 21 lines with a single T-DNA insertion locus. Even though all the lines showed a 3:1 Basta resistance segregation in the T2, we found three T3 lines in which there were multiple insertions in different chromosomes, suggesting that some of them were not expressing the transgenes properly. An interesting finding was that AtS/MAR10 40.01, the outlier line that showed abnormally high LUC expression, had two insertions very close to each other in chromosome 1, what could explain their behavior as a single locus in our segregation analysis and the increased transgene expression. The WGR data also allowed us to map the T-DNA insertion site of each line and to identify the deletions in the host genome associated with the insertion (Figure 3B and Table 1). Surprisingly, integration was not homogeneous among all chromosomes (we found none of the mapped insertions to be located in chromosome 2), and for Rb7 lines there was a clear preference for insertion within chromosome 3 (60%, 3 out of 5 lines) and with the T-DNA in the 3’->5’ direction (100%, 5 out of 5 lines), while for the rest of the lines chromosome 3 integrations and reverse T-DNA insertions only represented a 31% in each case (5 out of 16 for each) (Table 1).

**Figure 3.**
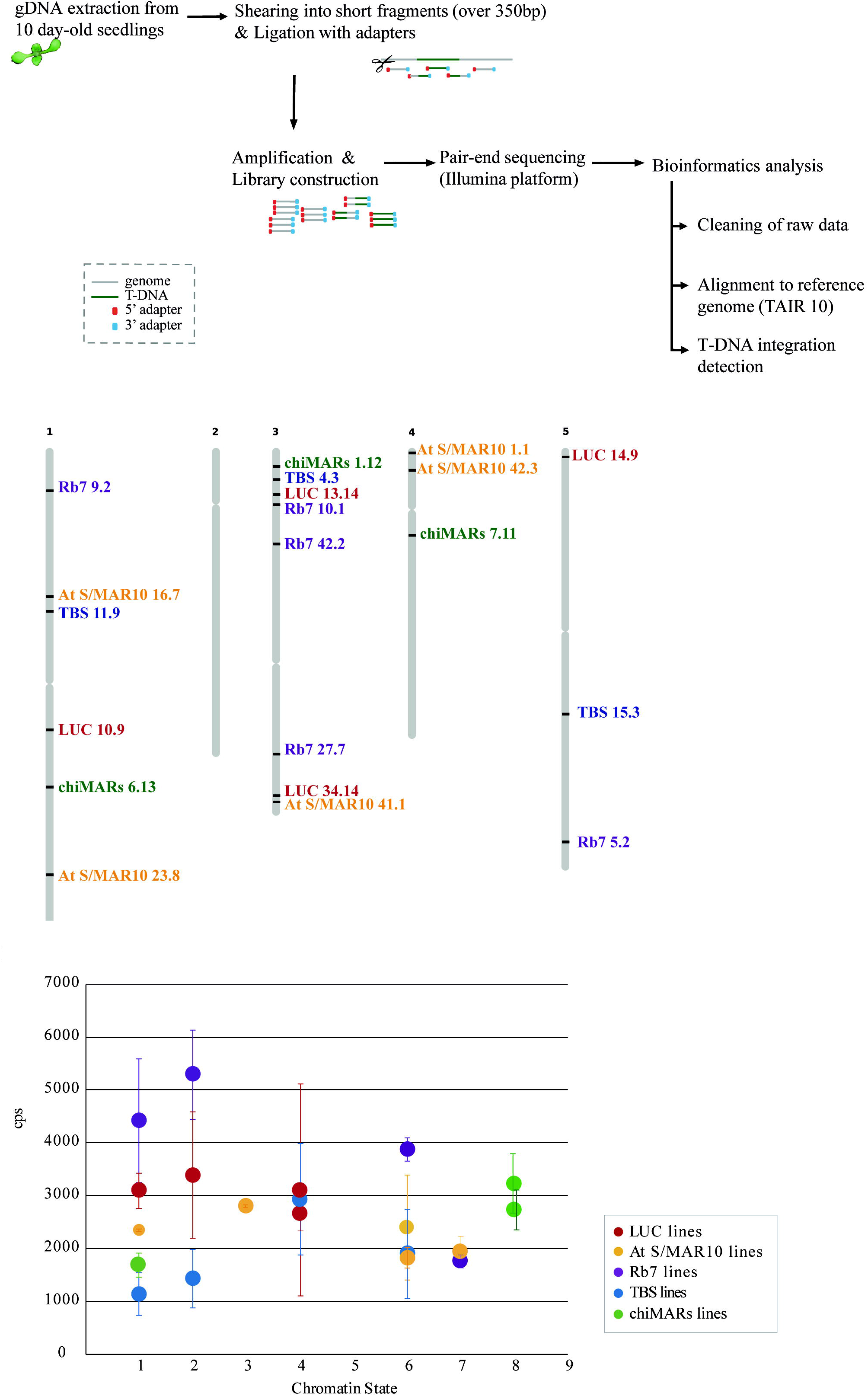
Analysis of insulator effect over T-DNA insertion. A) Scheme of the WGR pipeline B) Representation of the T-DNA insertion sites mapped within the five Arabidopsis chromosomes. C) Graph showing LUC activity versus chromatin state (Sequeira-Mendes et al., 2014) at T-DNA integration site. cps: counts per second.

**Table 1.**
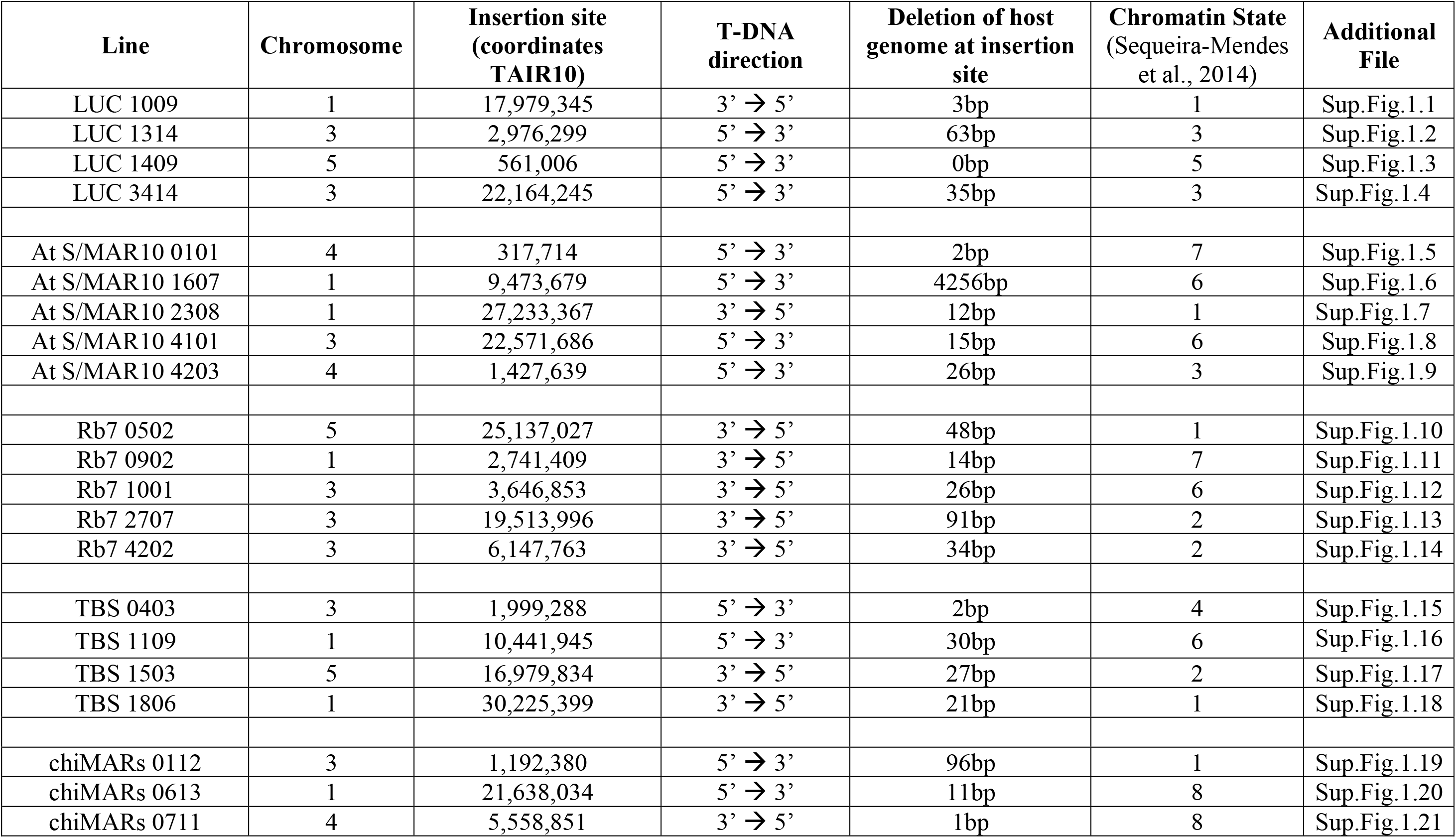
Details of the T-DNA insertions for single-copy lines based on WGR results.

The existence of a selection bias towards T-DNA integrations in euchromatin where the transgenes used for selection of transformants are expressed has been reported previously in the literature (Francis and Spiker, 2005). This was the case for most of the insertions we mapped (insertion sites in euchromatin, chromatin states 1 to 7 described in Sequeira-Mendes *et al*., 2014, Figure 3C), and when we plotted LUC activity versus state of the chromatin at the T-DNA insertion site, we could observe that lines grouped high or low depending on the construct they belonged to, and not left or right depending on the chromatin state where the T-DNA integration was located (Figure 3C). However, 2 lines carrying the chiMAR insulator presented T-DNA insertions in regions of the host genome featuring “chromatin state 8”, described as an A/T rich heterochromatic region characterized by methylated DNA and chromatin modifications such as H3K9me2 and H3K27me1 (Sequeira-Mendes et al., 2014).

We performed an analysis of the DNA methylation levels in the junction between the host genome and the T-DNA insertion for chiMAR line 6.13 and our results show that the DNA at the insertion site is indeed heavily methylated while the DNA of the T-DNA remains devoid of this chromatin modification even in the T3 generation, consistent with a boundary role of the insulator (Figure 4).

**Figure 4.**
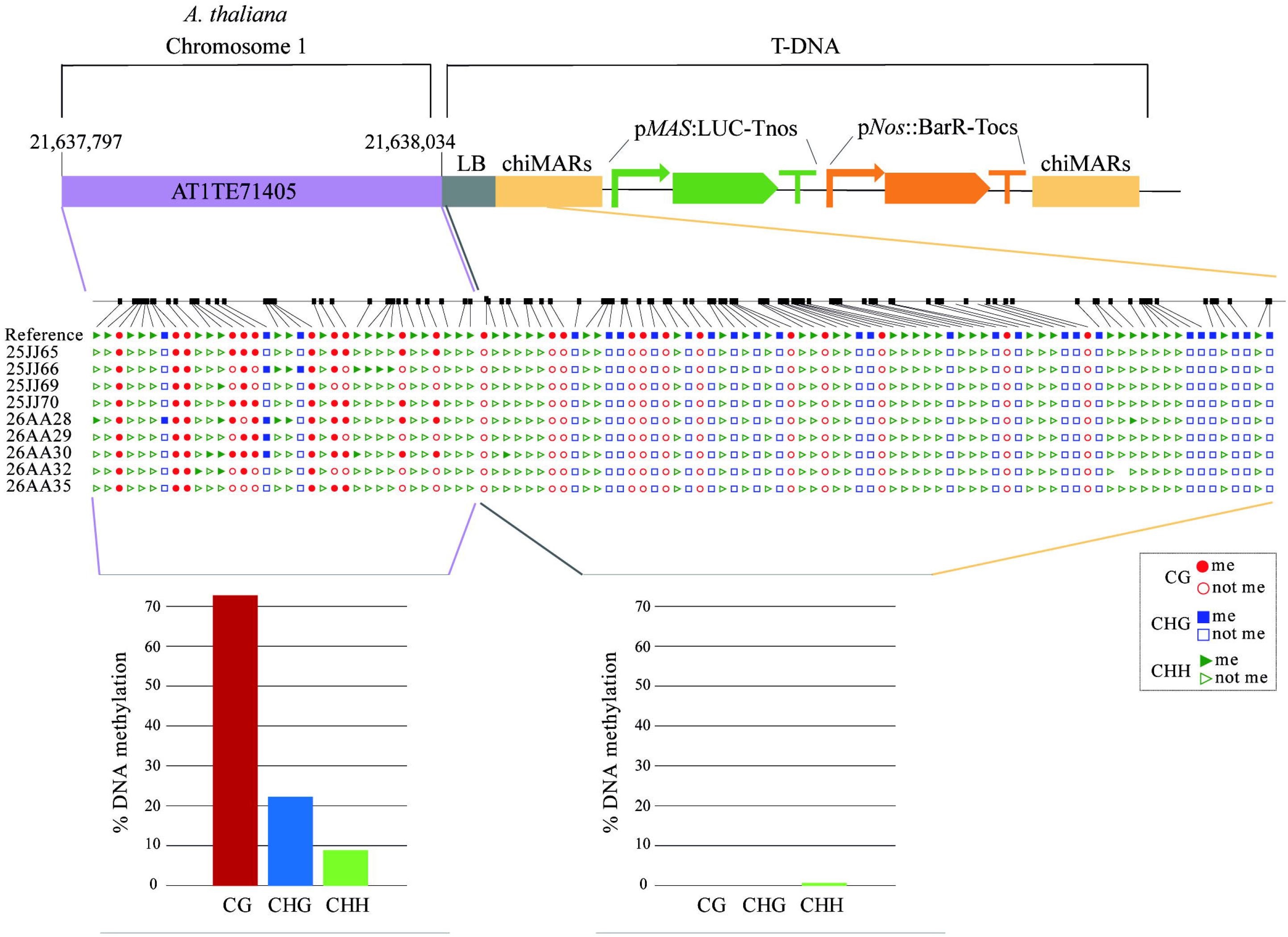
Analysis of the DNA methylation levels in the junction between the host genome and the T-DNA for line chiMAR 6.13. Upper part: schematic representation of the junction site. Middle part: graphical output of the methylation analysis (CyMate software) in 12 day-old seedlings of chiMARs 6.13 line. Red circles represent CG sites, blue squares represent CHG sites and green triangles represent CHH sites. Filled symbols indicate methylated cytosines while empty ones represent non methylated cytosines. Lower part: the graph shows the DNA methylation quantification of CG (red bars), CHG (blue bars) and CHH (green bars) cytosine contexts for the flanking sequence (left) and the T-DNA (right).

The data from WGR also allowed us to characterize the genomic sequence generated as a result of the T-DNA integration, and we could observe that for 8 out of 17 of the lines that contained insulator sequences, we had evidence of a lack of precision in the insertion of the RB, while that was not the case for any of the 4 control lines (Table 2). 3 out of 5 of the AtS/MAR10 lines contained vector backbone DNA (from outside the T-DNA region) integrated into the plant genome, while 3 out of 5 of the Rb7 lines, one AtS/MAR10 and one TBS line showed different degrees of truncation of the inserted T-DNA in the right border region. There was no evidence of truncation in the LB for any of the lines analyzed.

**Table 2.** Characterization of the genomic sequences generated as a result of the T-DNA integrations.

| Line | Chromosome | Insertion site<br>(coordinates TAIR10) | T-DNA direction | T-DNA 5' insertion site | T-DNA 3' insertion site |
| --- | --- | --- | --- | --- | --- |
| At S/MAR10 0101 | 4 | 317,714 | 5' → 3' | -35 (LB) | -959 (Tocs) |
| At S/MAR10 1607 | 1 | 9,473,679 | 5' → 3' | 0 (LB) | +3746 (RK2 TrfA) |
| At S/MAR10 4101 | 3 | 22,571,686 | 5' → 3' | -21 (LB) | +3825 (RK2 TrfA) |
| At S/MAR10 4203 | 4 | 1,427,639 | 5' → 3' | -17 (LB) | +1788 (pUC ori) |
| Rb7 0902 | 1 | 2,741,409 | 3' → 5' | -19 (LB) | -370 (Rb7) |
| Rb7 2707 | 3 | 19,513,996 | 3' → 5' | 0 (LB) | -515 (Rb7) |
| Rb7 4202 | 3 | 6,147,763 | 3' → 5' | -21 (LB) | -1052 (Rb7) |
| TBS 1503 | 5 | 16,979,834 | 3' → 5' | -5 (LB) | -2454 (Tocs) |

## Discussion

### Effect of insulators on transgene expression level and variation between lines

Most previous works have reported positive evidence of the effects of insulators on transgene expression, although some works can be found in the literature that report no such effect. The experiments were, however, very diverse in terms of species (some experiments had been done in tobacco and others in Arabidopsis) and in terms of method of transformation (some performed in primary transformants after regeneration and some in floral-dipped Arabidopsis).

It was an important motivation for this study to compare the effects of the different isolators in the same conditions: organism, developmental stage and transformation method. Our results do in fact support most results from literature, since we detect an increase in expression for lines where LUC is flanked by any of the four insulators, and previous negative results could reflect a dependency of the function of insulators on the experiment conditions.

Noteworthy, the use of AtS/MAR10, that had never been tested before for insulator activity, resulted in a moderate but very consistent increase in LUC expression.

In our hands, neither chiMAR, Rb7 nor TBS had an effect on reducing inter-line or inter-individual variation, in fact they increased them significantly. However, previous studies on the effect of chiMAR had highlighted its effect on the reduction of expression variability among transgenic lines (Mlynarova et al., 1995, 1994). This inconsistence could derive from a few factors in which our study differs basically from these other works. First, in our system we have used the pMAS promoter (versus the p35S used by Mlynarova et al. 1994 and Mlynarova et al. 1995) which never reaches such high levels of expression as the p35S, but that results in normally distributed expression levels in populations of transformants (De Bolle et al., 2003). It might be possible that the chiMAR works reducing the variance of strong promoters but its effect is not so apparent in promoters with an intrinsically low level of variation such as pMAS, like Mankin *et al*., 2003 described for Rb7. Second, in our study we have analyzed expression in homozygous T3 lines, that are already established lines with low variance in comparison with the T1 transformants analyzed by Mlynarova et al. 1994 and Mlynarova et al. 1995. It is interesting to note that the levels of variability between lines in the LUC control are in the same range as the variability between genetically identical individuals (around 30%), supporting the consistency and small intrinsic variance of our experimental set up in which we analyze T3.

In fact, it is striking that AtS/MAR10 is able to diminish inter-line variance, proving efficient in modifying both of the parameters measured, increasing transgene expression and reducing variability between lines, what makes it the best performing of the insulators analyzed.

### Effect of insulators on T-DNA insertion

Two interesting observations have been made regarding the effect of insulators on the insertion of T-DNAs. On the one hand, it is reported that T-DNA integrations recovered by selection are mostly located in “open chromatin” or euchromatin, while, without selection, integration is biased towards regions with marks of heterochromatin (Francis and Spiker, 2005). This is explained by the silencing of the selection genes when integration takes place within heterochromatin, a phenomenon that prevents transformant recovery. Our results show the ability of chiMAR to shelter T-DNAs from heterochromatin spreading and to allow for transgene expression regardless of the position effect.

On the other hand, the observation of an increased frequency of truncated T-DNAs in the lines containing insulators had been reported before by Li *et al*., 2008. Our results can be interpreted in the light of a role of insulators in the protection of transgenes at the right border end of the T-DNA from deletions. This would also explain the low correlation of expression between reporter genes located within the same T-DNA observed in many previous studies, and shown to improve by the use of insulators flanking them (Mlynarova et al., 1995). The preferential insertion of vector backbone in constructs harboring AtS/MAR10 cannot be explained by this rationale, though, and further experiments will be necessary to understand it.

As a general conclusion, we can state that there are many different insulators described in the literature with very different properties. Their functions might reflect differences in their action mechanisms and their use in transgenic constructs should depend on the needs of a specific experiment.

In our experimental setup, the best performing insulators were Rb7 in terms of increase of transgene expression, and AtS/MAR10 in terms of reducing variance.

Plant biologists should invest more efforts in the development of technologies that can render transgenes with high and stable expression with rapidity and ease. The future of synthetic biology and biotechnology projects depends on our ability to stabilize transgene expression and alleviate interference with the host genome regulation. In this work we show that the use of genetic insulators can help achieve these objectives with their simple addition at the flanks of the constructs used for transformation.

## Acknowledgements

Technical help from Cristina Vaca and Diana Coroian is greatly appreciated.

## Author contributions

AP and EC designed the experiments and analyzed the results. AP carried out the experiments. EC wrote the manuscript.

## Supplementary files 1

WGR data at the 21 mapped single insertion sites.

## List of Abbreviations

MAR: matrix attachment region
pNOS: nopaline synthase Agrobacterium gene promoter
TBS: transformation boost sequence
pMAS: mannopine synthase Agrobacterium gene promoter
S/MAR: scaffold/matrix attachment region
WGR: whole genome re-sequencing

